# A Mathematical Model Relating Pitocin Use During Labor with Offspring Autism Development in Terms of Oxytocin Receptor Desensitization in the Fetal Brain

**DOI:** 10.1101/446997

**Authors:** Mark M. Gottlieb

## Abstract

This paper develops a mathematical model describing the potential buildup of high oxytocin concentrations in the maternal circulation during labor in terms of continuous Pitocin infusion rate, half-life and maternal weight. Oxytocin override of the degradation of oxytocin by placental oxytocinase is introduced to model the potential transfer of oxytocin from the maternal circulation across the placenta into the fetal circulation, and from there into the brain of the fetus. The desensitization unit D equal to 1.8E6 (pg-min)/ml is employed to establish a desensitization threshold, and by extension; a down-regulation threshold as a function of oxytocin override concentration and continuous Pitocin infusion time, that could be a factor in the subsequent development of autism among offspring. Epidemiological studies by Duke University (S.G. Gregory, 2013), Yale University (O. Weisman, 2015) and Harvard University (A.S. Oberg, 2016) are discussed regarding Pitocin use and offspring autism development for an explanation of the weak correlations they identified. The findings of the Harvard epidemiological study are re-interpreted regarding Pitocin use, and its conclusion questioned. Further evaluations of the findings of these three epidemiological studies are called for to incorporate medical information on quantity of Pitocin used, continuous Pitocin infusion rate, length of labor and maternal weight to determine if a correlation can be established with offspring autism development above an empirically determined desensitization threshold for Pitocin use. Suggestions for research are discussed, including an alternative to continuous Pitocin infusion, pulsatile infusion of Pitocin during labor induction, that may mitigate possible offspring autism development.

**HIGHLIGHTS:**

- Buildup of oxytocin (OT) in the maternal circulation mathematically modeled.
- Relationship of OT half-life and OT concentration in maternal circulation identified.
- OTR Desensitization related to Pitocin infusion time and OT override concentration.
- Weak correlations for autism development in epidemiological studies explained.
- Examination called for of medical records of Pitocin use in epidemiological studies.

**OUTLINE OF PAPER:** Introduction

I. Development and application of mathematical model – Sections 1 through 6, notably: Figures 1 through 4
II. Discussion of oxytocin receptor desensitization as it pertains to mathematical model – Sections 7 and 8
III. Influence of mathematical model on interpretation of epidemiological studies – Sections 9 and 10
IV. Research considerations – Section 11, notably:Subsection 11.1 – Call for detailed epidemiological analysis of selected medical information indicated by mathematical model Subsection 11.2 – Review of pulsatile Pitocin infusion to mitigate possible offspring autism development indicated by mathematical model

## INTRODUCTION

An epidemiological study by the Duke University published in 2013, that was conducted in North Carolina found a modest association between labor induction and labor augmentation with the incidence of Autism among the offspring (S.G. Gregory, 2013) [1]. An epidemiological study in Denmark by Yale University published in 2015 found a mild association in males between use of Pitocin during labor augmentation and the incidence of autism among the offspring (O. Weisman, 2015) [2]. More recently, an epidemiological study by Harvard University published in 2016 investigated the association between labor induction and autism in Sweden and initially identified a weak correlation, (A.S. Oberg, 2016) [3]. However, using a novel statistical approach involving a family model relating the incidence of autism with labor induction among the offspring of mothers with multiple offspring, the Harvard study concluded that there was no correlation between labor induction, and by inference; the use of Pitocin and the incidence of autism among the offspring.

The objective of this paper is to provide a mathematical model relating the desensitization of oxytocin receptors (OTRs) in the brain of the fetus to the continuous infusion of Pitocin during labor, and by this mathematical model demonstrate that for long labors having long Pitocin infusion times with high Pitocin infusion rates desensitization of OTRs in the brain of the fetus may occur. Oxytocin receptor desensitization may be followed by down-regulation of oxytocin receptors [4], which was a concern of the Duke University study as a possible factor causing the development of autism in offspring [1]. This paper demonstrates that autism attributable to OTR desensitization arising from the use of Pitocin would be shown not to occur for labor inductions that have usual time-frames, which averages six hours [5], and may take as long as 12 hours or more for continuous Pitocin infusion [6].

However, this mathematical model does indicate that OTR desensitization may occur in some cases of long labor induction and long labor augmentation, in which high Pitocin infusion rates are administered over long labors with long Pitocin infusion times. OTR desensitization and subsequent down regulation may be associated with the development of autism [1]. Hence, the conclusion of the Harvard study may be in error, in that the outcome that they claim does not occur, no correlation of autism with labor induction, may occur for high infusion rates of Pitocin with long Pitocin infusion times during long labor inductions. Extrapolating data from these epidemiological studies to the population of the United States, it is possible to infer from the findings of this study that several hundred to one thousand or more offspring a year may be at risk of developing autism, following the use by their mothers of large quantities of Pitocin at high infusion rates during long labors. This comes to 0.5% to 1.0% of the number of cases of autism occurring each year in the United States.

This model was developed by the author to follow an earlier paper (M.M. Gottlieb, 2016) [7], in which it was shown that routine doses of Pitocin administered during typical labors would not result in OTR desensitization, and hence in autism attributable to OTR desensitization. However, the earlier paper did not consider the effects of high continuous infusion rates over the course of long labors with long Pitocin infusion times on OTR desensitization with possible offspring autism development, which this paper does.

## I. DEVELOPMENT AND EXPOSITION OF MATHEMATICAL MODEL (SECTIONS 1 – 6)

### 1. EXPLANATION OF FIGURE 1

To understand the buildup of Pitocin in the maternal circulation from a mathematical perspective, it is important to note that the concentration of Pitocin in the maternal circulation increases as a function of the half-life of Pitocin. The half-life of Pitocin in the maternal circulation is an indication of the relative efficiency of the liver and kidneys in removing Pitocin from the maternal circulation. Higher half-lives indicate lower Pitocin removal efficiency. The graph in Figure 1 indicates the relationship of the half-life of oxytocin in the blood with the percent oxytocin removal by the liver and kidneys and the percent oxytocin remaining in the blood after the removal. The indicated removal and retention of oxytocin is for a one-minute cycle of the blood circulation.

**Figure 1.**
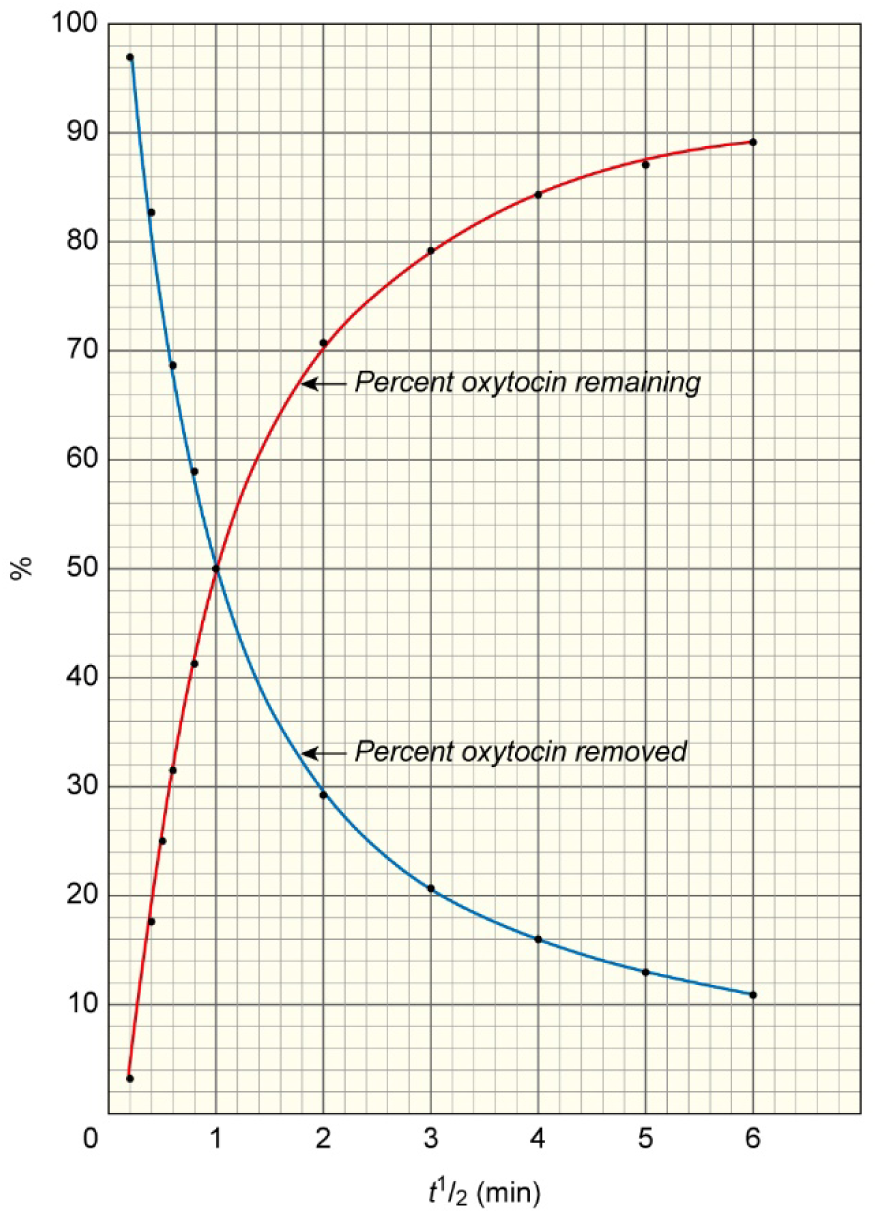
Percent Oxytocin Removal by Liver and Kidneys and Percent Oxytocin Remaining in Blood with One-Minute Circulation Time as a Function of Oxytocin Half-Life in the Blood.

### 2. BUILDUP OF PITOCIN CONCENTRATION IN THE MATERNAL CIRCULATION

In this section, I will offer a mathematical analysis to explain the appearance of concentrations of several hundred pg/ml or more of Pitocin in the maternal circulation, arising from Pitocin infusion into a vein in the uterus. The numbers follow arise out of an infusion rate of 20 mU/min, where 1 U or Unit of Solution, measuring one liter, introduced into the uterus contains 10 iU, or International Units of Pitocin with each iU containing 2 micrograms of Pitocin [8, 9]. The blood volume is 4.7 liters, which is for a 160-pound adult female, since the volume of blood for a woman is 65 ml/kg [10].

The average half-life of Pitocin in the maternal blood circulation decreases during pregnancy [11]. However, the half-life of OT in the maternal blood circulation may vary among women during labor, with less efficient removal of oxytocin by the kidneys and livers reflected in longer half-lives. The calculations are done for half-lives of 1 through 6 minutes, which appear in the literature for non-pregnant adults [12]. For this example, a half-life of 3 minutes was chosen. During late pregnancy, the half-life of oxytocin in the maternal circulation is reduced [12], though half-lives of 3 to 5 minutes have been reported [9, 11]. One minute is taken as the time needed to complete one cycle of the maternal blood circulation.

It is important to note that the concentration of Pitocin from continuous Pitocin infusion will increase until the rate at which it is added to the maternal circulation equals the rate at which it is removed by the kidneys and liver [13].

To illustrate the calculations used in this section, an amount of Pitocin is added for each one-minute cycle of 20 mU/min, which is a typical value for the continuous infusion of Pitocin during labor. Note, that the infusion of 20 mU/min for one minute corresponds to the addition of 4E5 pg of OT, when 10 iU of Pitocin is introduced into a liter of solution referred to as a Unit U, and 1 iU of Pitocin equals 2 micrograms of Pitocin [12,14]. Since this is in 4.7 liters of blood, the concentration is 85 pg/ml that is increased during each cycle. However, there is also the removal rate of the Pitocin, which is expressed as an exponential term Exp((−ln2/t1/2)t’), where t1/2 is the half-life for the Pitocin in the maternal circulation, ln2 is the natural logarithm of 2 equal to 0.6931, and t’ = 1 minute is the time for one cycle in the circulation of the maternal blood.

Hence, the concentration of the Pitocin at the end of a one-minute cycle is the product of 85 pg/ml, the amount of Pitocin added in one cycle, and the exponential removal rate over one cycle, which is Exp(−ln2/t1/2) for the concentration of OT after the first cycle. This is equal to 85Exp(−ln2/t1/2) pg/ml. For the next cycle, another 85 pg/ml increase in concentration takes place, but there is the removal of some of the Pitocin remaining from the first cycle plus some of the Pitocin from the second cycle. This can be expressed as the concentration C for two cycles, where…

C =85[Exp((−ln2/t1/2)(t’)) + Exp2((−ln2/t1/2)(t’))] pg/ml

This process continues and can be expressed in general terms as:

C = 85ΣExp((i)(−ln2/t1/2)(t’)) pg/ml

For the buildup concentration C of Pitocin, where i runs from 1 to some large value n. Each hour Pitocin is infused n = 60, assuming a one-minute maternal blood circulation rate. In mathematics, this is known as a geometric series [15]. The sum C of a geometric series, where n is large and i runs from 0 to n, is given by:

C = k[1/(1-r)], where r is equal to Exp(−ln2/t1/2), which is the ratio of successive terms in the geometric series and is less than 1. In this example

k = [(IR)(10E-3)(10)(2E6)]/V pg/ml = [(IR)(2E4)/V] pg/ml

where k is the concentration in pg/ml of the Pitocin that is added to the maternal circulation per minute, IR is the continuous infusion rate in mU/min and V is the maternal blood volume, which is equal to the maternal weight W in kg times (65 ml/kg) [10]. In our case, IR = 20 mU/min and the volume V = 4.7E3 ml for a maternal weight of 160 pounds. Therefore, k = 85 pg/ml in a one-minute cycle, as indicated above. However, since Pitocin is removed during the first cycle, i runs from 1 to n, the formula becomes:

C = k[1/(1-r) - 1] = k[r/(1-r)] pg/ml

Where, r = Exp(−ln2/t1/2) and the buildup concentration of oxytocin C = k[Exp(−ln2/t1/2)/(1-Exp(−ln2/t1/2))] pg/ml. The term in the brackets [ ] is referred to as a Multiplier M. For the half-lives considered in this paper of 1 to 6 minutes, the concentration C comes close to the limit r/(1-r) within 20 or 30 minutes, as it continues to increase with the continuous infusion of Pitocin. For a value of t1/2 = 3 minutes, V = 4.7 liters and k = 85 pg/ml, the concentration C = 85(0.794/(1-0.794)) pg/ml = 85(3.85) pg/ml = 327 pg/ml for the buildup concentration of OT in the maternal blood circulation for an infusion rate of 20 mU/min with a maternal blood circulation rate of one-minute. See Figure 2 below for an illustration of this buildup process by the summation of each succeeding exponential term.

**Figure 2.**
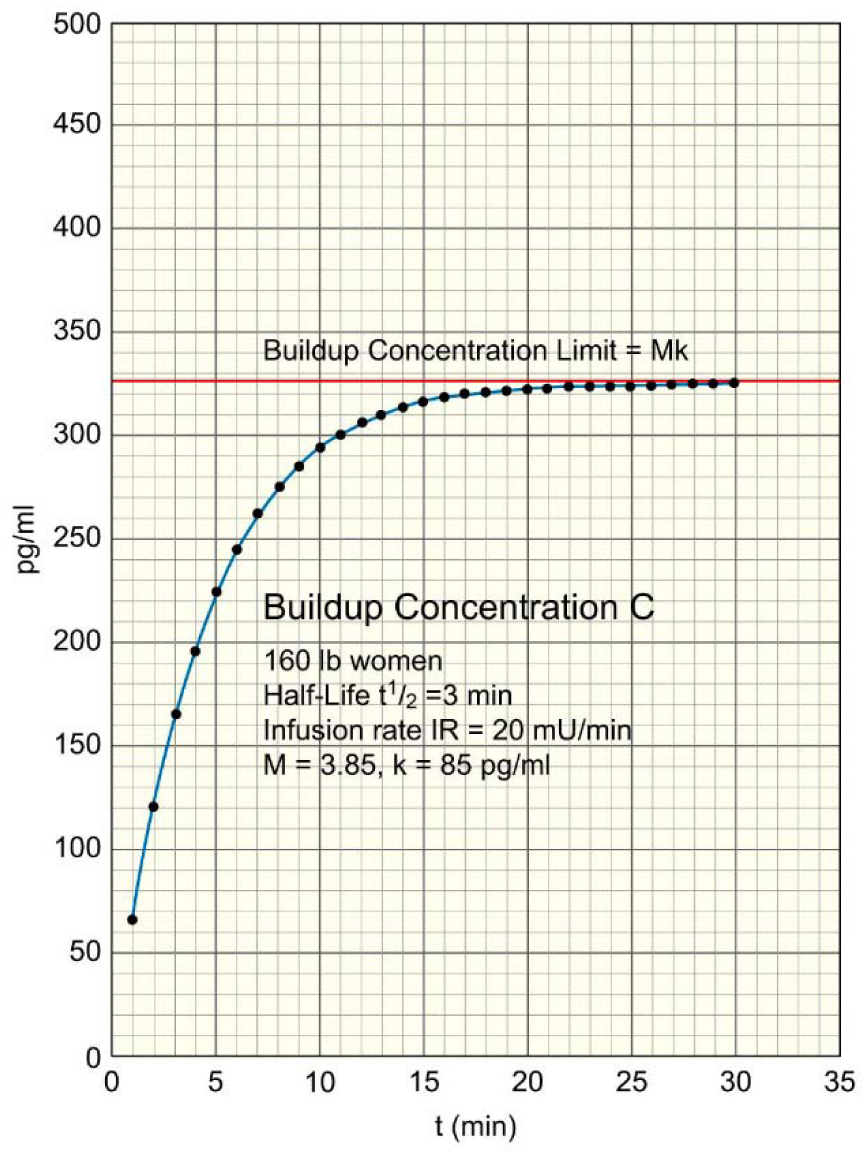
Buildup of the baseline Concentration C of Pitocin in the Maternal Circulation from Continuous Pitocin Infusion. Note the parameters of maternal weight, infusion rate, and half-life, and the values of the Multiplier M and k, which are determined by the half-life and infusion rate in a one-minute blood circulation cycle as indicated in Tables 1 and 2 in Section 2.

The term r/(1-r) is referred to as a Multiplier, M, and is unique to each half-life considered. Note that the buildup concentration C is equal to Mk, or C = Mk. The values for M for the half-lives considered, and the values for k for the buildup concentrations of Pitocin in one cycle that arise from the indicated Pitocin infusion rates, IR, for maternal weights of 160 pounds and 128 pounds in a 1-minute blood circulation cycle are given in the Table 1 and Table 2, below.

**Table 1.** Values of the Multiplier M for the indicated half-lives.

| Half-life (min) | M |
| --- | --- |
| 1 | 1.00 |
| 2 | 2.41 |
| 3 | 3.85 |
| 4 | 5.29 |
| 5 | 6.73 |
| 6 | 8.17 |

**Table 2.**
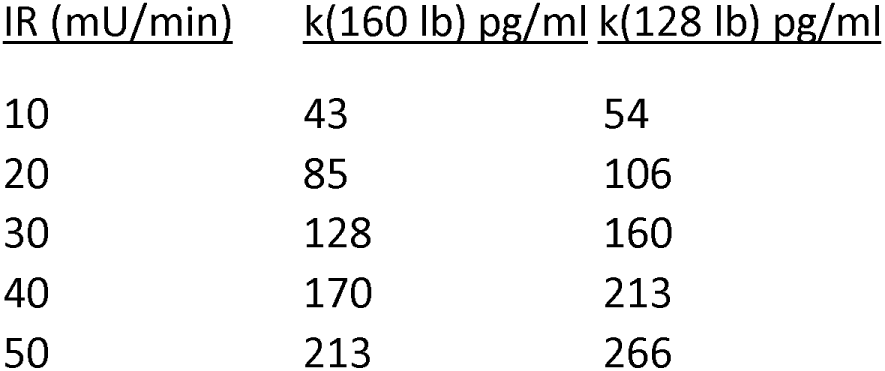
Values of k for the concentration of Pitocin that is added in 1-minute for the indicated Pitocin infusion rates, IR, and the indicated maternal weights.

| IR (mU/min) | k(160 lb) pg/ml | k(128 lb) pg/ml |
| --- | --- | --- |
| 10 | 43 | 54 |
| 20 | 85 | 106 |
| 30 | 128 | 160 |
| 40 | 170 | 213 |
| 50 | 213 | 266 |

### 3. EXPLANATION OF FIGURE 2

Figure 2 indicates the incremental buildup of the concentration of oxytocin in the maternal circulation as the summation of succeeding exponential terms indicating the remaining oxytocin after each 1-minute cycle. Note that this buildup has an upper limit expressed as Mk (see Tables 1 and 2 in Section 2), which is asymptotically approached. The 30-minute value of the concentration is within 0.1% of the buildup limit Mk and is stable. This 30-minute period is the usual length of time that Pitocin is kept at each Buildup interval until the buildup of the necessary Pitocin concentration during continuous Pitocin infusion is reached, which is accepted medical practice [12,14].

In actual practice, the buildup of the concentration of Pitocin from continuous Pitocin infusion is done in a very gradual manner from one buildup concentration to the next as the continuous Pitocin infusion level is increased. The characteristics of the buildup process for each individual buildup concentration of Pitocin at each specific continuous Pitocin infusion rate are more complex than indicated in Figure 2. There is an increase each minute for each succeeding incremental concentration from the prior concentration, and then a decrease, until the stable concentration level C = Mk is reached when each increase is followed by a nearly equal decrease each minute before the next incremental concentration is reached. However, the graph in Figure 2 correctly indicates the characteristics and baseline levels of the resultant buildup concentration C = Mk.

The more conservative approach for indicating the lower baseline value of the buildup concentration indicated in Figure 2 was chosen to facilitate the mathematical modeling of the buildup concentration, since otherwise; the actual value of the buildup concentration at any given time would be uncertain because of the nature of the fluctuations resulting from the increase and decrease of the concentration over each 1-minute cycle. The incremental buildup concentrations shown in Figure 2 below are for their baseline levels.

### 4. EXPLANATION OF FIGURE 3

This graph in Figure 3 depicts a linear relationship for each Pitocin infusion rate for the buildup concentration of Pitocin in the blood in the maternal circulation plotted against the half-life of Pitocin in the maternal circulation for mothers with weights of 160 pounds and 128 pounds, respectively. The relationship between C and t1/2 over values of t1/2 from 1 minute to 6 minutes for Pitocin infusion rates of 10 mU/min to 50 mU/min is given in Figure 3 for women weighing 160 pounds with a blood volume of 4.7 liters on the left, and for smaller women weighing 128 pounds, 20% less, with a volume of 3.8 liters of blood on the right. The linear relationship of each infusion rate with half-life should be noted.

**Figure 3.**
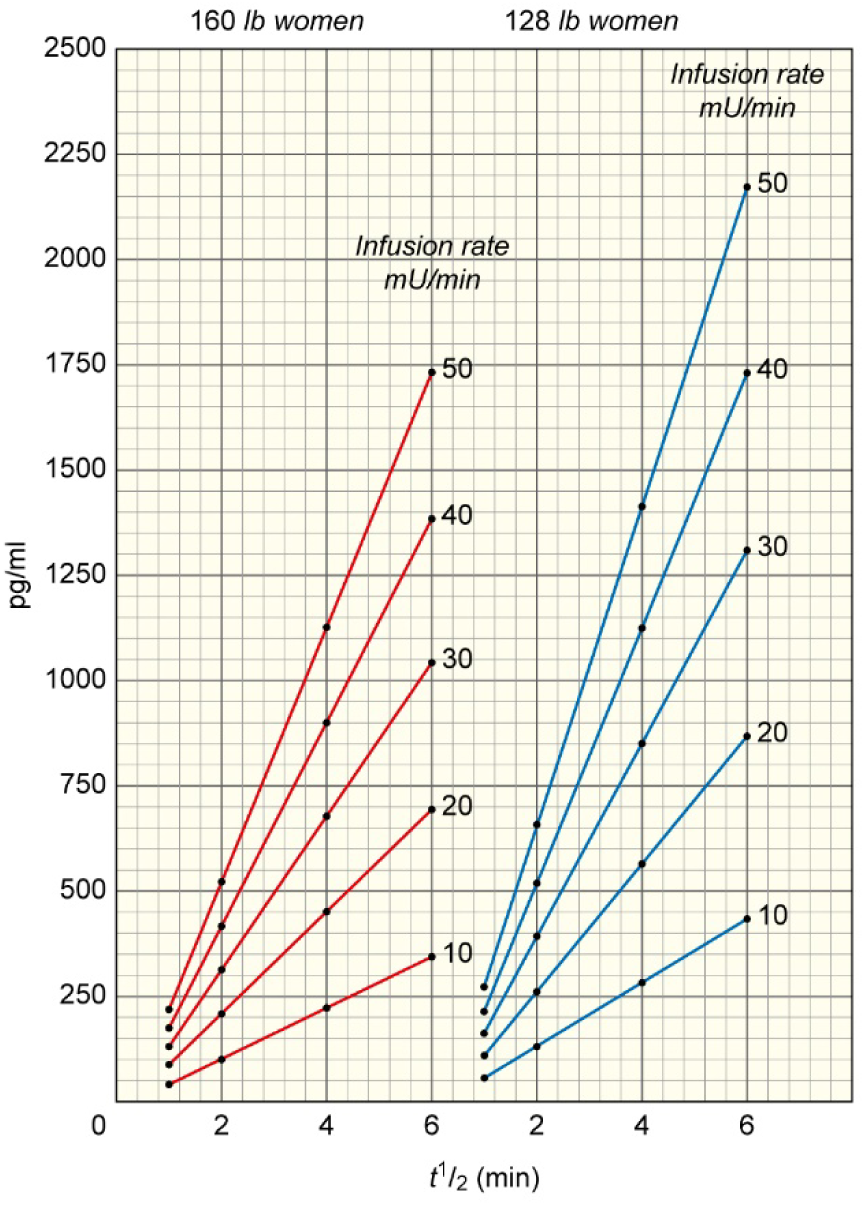
Oxytocin baseline Buildup Concentrations in Maternal Circulation in terms of Pitocin Infusion Rate, Oxytocin Half-Life and Maternal Weight as indicated in Tables 1 and 2 in Section 2.

The values of the continuous Pitocin infusion rates are taken from the values of M and k in Tables 1 and 2 in Section 2, where the baseline Buildup Concentration C = Mk for each combination of M and k. Note, that the value of the half-lives are directly related to the values of M, and the values of the continuous Pitocin infusion rates and maternal weights are directly related to the values of k. Mothers with smaller weight will have higher concentrations of Pitocin in the maternal circulation for a given infusion rate than larger mothers, in that blood volume is proportional to the mother’s weight, and the same quantity of Pitocin introduced into a smaller blood volume will have a higher concentration that is inversely proportional to the maternal blood volume.

The significant influence of the half-life of oxytocin in the maternal circulation on the buildup concentration of oxytocin in the maternal circulation, as indicated in Figure 3, may not have been previously realized. Its indication by the mathematical model developed in this paper follows from the buildup of the Pitocin concentration indicated in Figure 2. The variations in the efficiency of the maternal liver and kidneys in removing oxytocin from the maternal circulation are related to the half-life of the Pitocin in the maternal circulation, as indicated in Figure 1. Lower efficiencies of oxytocin removal are reflected in higher half-lives, and consequently higher oxytocin buildup concentrations in the maternal circulation, as indicated in Figure 3.

One observation arising from this graph is that higher oxytocin concentrations in the maternal circulation are more likely to result in oxytocin diffusing through the placenta.

It should be noted, as mentioned Section 2, the half-life is shorter during late pregnancy, though values of 3 to 5 minutes have been reported. However, it is conceivable that some mothers with very inefficient kidneys and livers may have longer half-lives for oxytocin removal. It should also be noted that for mothers with higher Pitocin infusion rates and higher half-lives, Figure 3 indicates that the concentrations of Pitocin in the maternal blood circulation can be several times higher than the concentrations for mothers with lower half-lives and lower Pitocin infusion rates.

### 5. COMPOSITE DESENSITIZATION UNIT “D”

In the author’s earlier paper (M.M. Gottlieb, 2016) [7], a composite unit “D ” equal to 1.8E6 (pg-min)/ml, relating the product of oxytocin concentration and oxytocin exposure time for the OTRs, was established to describe the desensitization threshold for the OTRs. Studies of OTR desensitization as a function of oxytocin concentration and exposure time are scarce. One study was identified (C. Robinson, 2003) [16]. From the information in this study, it was possible to develop the value of D by multiplying the numbers for the concentration of Pitocin, 1E4 pg/ml, and the time, 1.8E2 min, needed to achieve desensitization, which equals 1.8E6 (pg/ml)(min), and is expressed in terms of the composite unit as 1.8E6 (pg-min)/ml.

This establishes a relationship between oxytocin concentration and the time T for desensitization using the value of D, the threshold desensitization unit, as a constant used over a wide range of oxytocin concentrations and oxytocin exposure times. In other words, the oxytocin concentration C = D/T, where the concentration C is in pg/ml and the time T is in minutes and D = 1.8E6 (pg-min)/ml.

Knowing the average infusion rate in mU/min and the length of time at that infusion rate, it becomes possible using the model mentioned above to indicate the possibility of desensitization to occur. To do this, the concept of an upper limit of the ability of placental oxytocinase to degrade oxytocin is introduced. The concentration of oxytocin from the maternal circulation that exceeds it results in a smaller concentration of oxytocin that diffuses across the placenta, which is called the oxytocin override concentration. If this smaller oxytocin override concentration is in the range indicated in Figure 4 for OTR desensitization to occur, then the length of time at this smaller concentration the OTRs in the fetal brain are exposed permit calculations of desensitization of the OTRs using the equations above, C = D/T. The plausibility of this phenomenon arises from the development of the mathematical model in this paper that depicts the buildup of high oxytocin concentrations in the maternal circulation indicated in Figures 2 and 3, and the use of the desensitization unit D for the calculations indicated in Figure 4.

**Figure 4.**
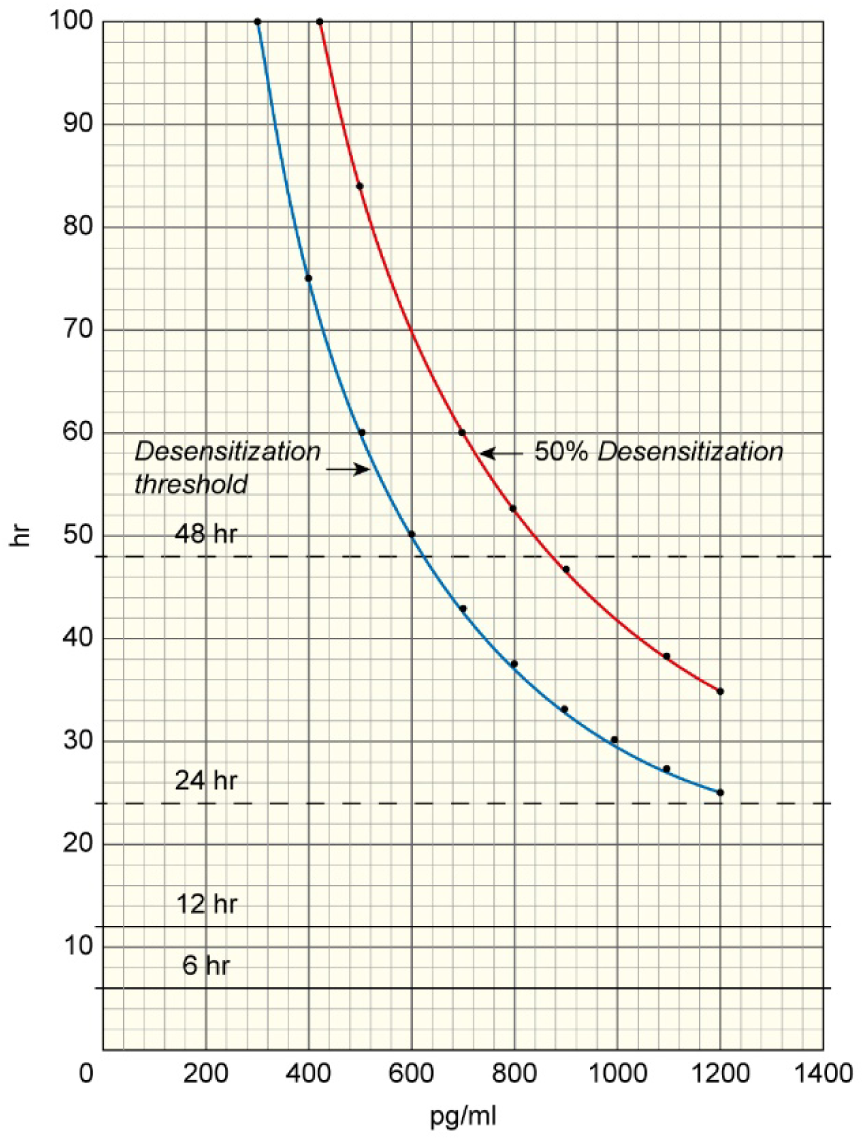
Oxytocin Override Desensitization Threshold and 50% Desensitization Level as Functions of Pitocin infusion time T during labor and Oxytocin Override Concentration C, where CT = D = 1.8E6 (pg-min)/ml for the Oxytocin Override Desensitization Threshold.

### 6. EXPLANATION OF FIGURE 4

Oxytocinase, which is an enzyme that degrades oxytocin, is found in high concentrations in the placenta during labor [17]. The contention in this paper is that very high concentrations of Pitocin in the maternal circulation in the placenta overwhelm the ability of the oxytocinase in the placenta to degrade all the Pitocin in the placenta from the maternal circulation. Consequently, some of the Pitocin may diffuse across the placenta and enter the fetal circulation (A. Malek, 1996) [18]. This phenomenon is referred to in this paper as “Oxytocin Override” of placental oxytocinase degradation of oxytocin.

As shown in Figure 3, the infusion of Pitocin into a vein in the uterus during labor can greatly increase the concentration of Pitocin in the maternal circulation. The mathematical model developed in this paper assumes nearly all the Pitocin introduced into the vein in the uterus mixes with the blood in the maternal circulation, some of which goes into the placenta.

The buildup of Pitocin in the maternal circulation has been discussed in the literature in terms of its effect on oxytocin concentration in the fetal circulation (C. Patient, 1999) [19]. However, from the analysis in that paper [19], the concentrations of oxytocin indicated in the maternal circulation may not have been sufficient to achieve oxytocin override of placental oxytocinase degradation of oxytocin, and thus would not enter the fetal circulation. This is noteworthy, in that the effect of maternal weight and oxytocin removal half-life may not have been incorporated into the research considerations that determined the findings of that paper.

Oxytocin override concentrations and corresponding desensitization times are given by the following relationship:

D/C(override) = T, where D = the desensitization constant of 1.8E6 (pg-min)/ml developed from the literature (M.M. Gottlieb, 2016), (C. Robinson, 2003) [7,16], C(override) = oxytocin override concentration, and T = the desensitization time, the time needed to reach the desensitization threshold at the specified concentration in the fetal circulation equal to C(override).

This relationship is indicated in the graph in Figure 4 for the desensitization threshold and for 50% desensitization over a range of oxytocin override concentrations and desensitization times, in which the 50% desensitization increase requires 40% longer desensitization time, is indicated in the literature [16]. It is apparent from Figure 4 that OTR desensitization would require very high oxytocin override concentrations to occur during the usual time frames for labor induction. Labor induction typically occurs in the first 6 hours of labor [5], the average time for labor induction, which is indicated by the 6hour line on the graph for continuous Pitocin infusion time, and many women may need 12 hours or more of Pitocin infusion [6], indicated by the 12-hour line on the graph in Figure 4. These time-frames would require very high oxytocin override concentrations that may possibly exceed the oxytocin buildup concentrations in the maternal circulation indicated in Figure 3 and would therefore be impossible to achieve.

However, Figure 4 indicates that longer desensitization times are consistent with lower oxytocin override concentrations, taking-into-account the use of oxytocin for both labor induction and labor augmentation. For long labors, indicated by the lines for 24 to 48 hours of Pitocin infusion time in Figure 4, the indicated oxytocin override concentrations in Figure 4 are less than the high concentrations of Pitocin in the maternal circulation indicated in Figure 3 at the higher continuous Pitocin infusion rates and half-lives and are therefore plausible. As indicated in Figure 3, high Pitocin infusion rates, longer half-lives and mothers with smaller weights serve to increase the concentrations of Pitocin in the maternal circulation, making oxytocin override concentrations high enough to achieve desensitization during long labors more likely.

## II. DISCUSSION OF OXYTOCIN RECEPTOR DESENSITIZATION (SECTIONS 7 AND 8)

### 7. OXYTOCIN OVERRIDE AND OXYTOCIN RECEPTOR DESENSITIZATION

The concentration of Pitocin in the fetal circulation may be high enough to result in desensitization of OTRs in the brain of the fetus, if the infusion rate of the Pitocin is high over the course of long labors with long Pitocin infusion times. As shown in Figure 4, the relationship between the concentration of Pitocin, C, and the desensitization unit D, where D = CT = 1.8E6 (pg-min)/ml, may be applied to the concentrations of Pitocin in the cerebral spinal fluid.

It is noteworthy that the blood-brain barrier of the fetus may be permeable to oxytocin [20]. For the purposes of this study, no inhibition of oxytocin crossing the blood-brain barrier in the fetus has been assumed. Pitocin may cross the blood-brain barrier in the fetus in high concentrations, resulting in high concentrations in the cerebral spinal fluid in the brain. It is possible to model the buildup of oxytocin in the fetal brain in terms of oxytocin override concentrations and desensitization times for the desensitization of OTRs in the fetal brain from information in Figure 4. This is so, since the oxytocin concentration in the fetal brain would most likely be equal or close to the oxytocin override concentration that diffuses across the placenta and enters the fetal circulation, since no inhibition of oxytocin transfer across the blood brain barrier has been assumed.

For high enough concentrations of Pitocin in the maternal circulation resulting from high rates of Pitocin infusion, the concentration of Pitocin that may enter the fetal circulation may be high. However, if it is not high enough, or is sufficiently high but not for long enough, OTR desensitization will not occur. Also, it is important to note that long Pitocin infusion times imply long labors, however; long labors do not necessarily imply long Pitocin infusion times, as alternative procedures and the stopping of Pitocin infusion may be done. It is noteworthy that oxytocinase does not leak into the fetal circulation [21]. Consequently, there is no degradation of oxytocin in the fetal circulation attributable to oxytocinase in the fetal circulation. While it is known that the fetal secretion rate of oxytocin increases significantly during labor with oxytocin concentrations of 73 pg/ml measured in the umbilical artery [22], this paper does not consider the endogenous production of oxytocin by the fetus in its calculations of oxytocin concentrations in the fetal circulation. Likewise, this paper does not consider the endogenous production of oxytocin in the maternal circulation in its calculations of oxytocin override concentrations [22].

In this regard, one study indicated very high levels of endogenous oxytocin in the maternal circulation with no contribution from Pitocin, when Pitocin infusion was performed during labor (M. Prevost, 2014) [23]. However, this study indicated that oxytocin was not extracted from the samples of plasma, which underwent enzyme limited immunoassays to measure the oxytocin levels. In a separate study on oxytocin measurement, it was found that oxytocin samples that underwent enzyme limited immunoassay without extraction of the oxytocin from the plasma could lead to measurements that were a factor of a hundred or more in error (A. Szeto, 2011) [24]. Therefore, the findings indicated in the first study [23] are considered erroneous.

The action of oxytocinase during pregnancy has been examined in the literature [25,26]. It is realistic to presume that there is an upper limit to the ability of placental oxytocinase to degrade all the oxytocin entering the placenta from the maternal circulation. While oxytocin override has not been confirmed by measurements, it is plausible that the higher the concentration of oxytocin in the maternal circulation, the more likely oxytocin override is to occur.

Earlier studies of Pitocin infusion, showing that the transport of oxytocin from the maternal circulation across the placenta into the fetal circulation does not occur (C. Patient, 1999), (J. Sjoholm, 1969) [19, 26], may not have considered the cases indicated in this paper for high infusion rates for smaller women with kidneys and livers that are less efficient at removing oxytocin, which would be reflected in longer half-lives and higher oxytocin concentrations, as indicated in Figure 3. The oxytocin concentrations in the maternal circulation in these studies [19,26] may not have exceeded the limit of placental oxytocinase to degrade the oxytocin.

### 8. FACTORS AFFECTING OXYTOCIN RECEPTOR DESENSITIZATION DURING LABOR IN THE FETAL BRAIN

The following factors may affect the concentration of Pitocin during labor and the subsequent desensitization of OTRs in the fetal brain:

8.1 The initial factor relating the use of Pitocin during labor with OTR desensitization and the possible development of autism among some of the offspring is the total quantity or units of Pitocin used, since this relates directly with the Pitocin infusion rate and the Pitocin infusion time.

8.2 A second factor relating the use of Pitocin during labor with OTR desensitization and the possible development of autism among some of the offspring is the continuous Pitocin infusion rate in mU/min [14] during the labor, where a unit U of Pitocin is 10 iU (International Units of Pitocin, where 1 iU = 2 ug of Pitocin) in 1 liter of solution. The higher the infusion rate of Pitocin the higher the potential oxytocin override and subsequent OTR desensitization.

8.3 It can be shown mathematically that high infusion rates of Pitocin may result in a disproportionately high concentration of Pitocin in the fetal circulation, relative to the same quantity of Pitocin introduced at lower infusion rates. The placenta can only produce a limited amount of oxytocinase to degrade the Pitocin from the maternal circulation. Once the maximum ability of the oxytocinase to degrade the Pitocin is reached, additional Pitocin may get though with more getting through for higher infusion rates than for the same quantity of Pitocin used in lower infusion rates. The implication from the discussion in this paper for the use of Pitocin during long labors with long Pitocin infusion times at high Pitocin infusion rates is that, where possible; Pitocin should be used at lower infusion rates, since that may limit the possible development of autism in the offspring.

8.4 The concentration and quantity of oxytocinase in the placenta may decrease during labor by means of leakage from the maternal circulation in the placenta into the maternal blood circulation as-a-whole [25]. A lower concentration of oxytocinase in the maternal circulation in the placenta may result and lead to greater oxytocin override, since the upper limit of Pitocin degradation by oxytocinase in the placenta would be lower. The concentration of oxytocinase in the maternal circulation as-a-whole would increase, increasing the degradation of oxytocin in the maternal circulation, however; the degradation of oxytocin in the maternal circulation by oxytocinase is small [26]. It can be shown mathematically that leakage of oxytocinase from the placenta to the blood would not compensate for a lower concentration of oxytocinase in the placenta and the reduction of its upper limit to degrade oxytocin. Consequently, due to the leakage of oxytocinase from the placenta, the placental oxytocinase limit for the degradation of oxytocin may decline and the oxytocin override concentration may increase. This may result in a greater likelihood and degree of desensitization of OTRs in the brain of the fetus. Placentas may vary in the concentration and quantity of oxytocinase they produce, and in the extent of their leakage of oxytocinase and the time over which it occurs.

8.5 Another factor is the potential increase in the concentration of oxytocin in the cerebral spinal fluid, resulting from the addition of Pitocin to the cerebral spinal fluid that crosses the blood brain barrier. The added concentration of Pitocin in the cerebral spinal fluid may itself lead to Pitocin’s action as a neurotransmitter and neuromodulator acting on magnocellular neutrons in the hypothalamus that produce the oxytocin. This may cause the production of even more oxytocin, which may result in a further increase in the concentration of oxytocin in the cerebral spinal fluid. This could be a factor affecting the desensitization of the OTRs and may cause OTR desensitization to occur at lower oxytocin override levels.

8.6 Another factor affecting the desensitization of OTRs by Pitocin is the half-life of the Pitocin in the maternal blood circulation, which is directly related to the efficiency of the kidneys and liver of the mother in removing oxytocin. As shown in Figures 2 and 3, the concentration of Pitocin in the maternal circulation is strongly influenced by the half-life of the Pitocin in the maternal circulation. Higher half-lives may result in significantly higher concentrations of Pitocin in the maternal circulation, and hence; greater oxytocin override, which may result in greater desensitization of OTRs in the brain of the fetus.

8.7 Another factor affecting the concentration of Pitocin entering the cerebral spinal fluid that may cause OTR desensitization in the brain of the fetus is the blood volume of the maternal circulation, in that for the same Pitocin infusion rate, a smaller blood volume in the maternal circulation will have a higher concentration of Pitocin as indicated in Figure 3, since blood volume and Pitocin concentration are inversely proportional. Higher Pitocin concentrations may result in greater oxytocin override, and hence; greater desensitization of OTRs in the brain of the fetus.

8.8 The fluctuations of the buildup concentrations mentioned in Section 4 that increase the magnitude of these buildup concentrations above the baseline may result in fluctuations in the oxytocin override concentrations that are indicated in Figure 4. These fluctuations may vary as a function of both the continuous Pitocin infusion rate and the half-life of the Pitocin in the maternal circulation. These fluctuations may result in a reduction of the time during labor that is needed for OTR desensitization in the brain of the fetus that is indicated in Figure 4.

8.9 The presence and concentration of OTRs in the brain of the fetus at term bears discussion. It has been found that just before birth estradiol has a very important role in influencing the number and concentration of OTRs in the brain of the fetus, increasing them by as much as a factor of five (M. Soloff, 1983) [27]. Variation in the ambient concentration of estradiol may significantly influence the concentration of oxytocin receptors in the brain of the fetus, which may be a factor in the onset of autism [28]. High oxytocin concentrations in the cerebral spinal fluid may interfere with the development of new OTRs in the brain of the fetus, as they do for existing OTRs during desensitization and down-regulation. This, in addition to the reduction in the numbers of OTRs that may result from lower levels of estradiol [27], may cause the possible development of autism to be more severe. This is so, because lower levels of estradiol may serve to augment the desensitizing action of oxytocin override in reducing the number of OTRs.

8.10 It has been found in a study by Stanford University, that that genetic variability in the OTR gene may be a risk for the development of autism (K.J. Parker, 2014) [29]. It is conceivable, that because of the genetic variability of the OTR gene, the OTRs of autistic children may be more vulnerable to desensitization, and consequently; undergo desensitization in a manner described by a smaller desensitization Unit D. Consequently, there may be less time needed for desensitization for any given oxytocin override concentration, or a lower oxytocin override concentration for any given desensitization time for fetuses having the variability in the OTR gene described in the Stanford study. The effect, if any, of the genetic variability of OTRs on their vulnerability to desensitization by oxytocin was not addressed in the Stanford study. Likewise, it is conceivable that these OTRs may not respond to the presence of estradiol during labor in the same way as the normal OTRs indicated in Subsection 8.9. The effect if any of the influence of estradiol on the development of the OTRs with genetic variability, as discussed for normal OTRs in Subsection 8.9, was also not discussed in the Stanford study.

8.11 Fetal production of oxytocin [22] may conceivably occur with the presence of oxytocin override concentrations entering the fetal circulation. This would increase the concentration of oxytocin in the fetal circulation beyond the oxytocin override levels, and consequently; lower the time-frames for desensitization of OTRs in the fetal brain or increase the likelihood or degree of this desensitization for a given time-frame. Similarly, endogenous oxytocin in the maternal circulation could serve to increase the oxytocin override concentration and its impact on desensitization of the OTRs in the fetal brain [22].

8.12 The desensitization unit D, which is treated as a constant equal to 1.8E6 (pg-min)/ml throughout this paper may vary over the ranges of Pitocin concentrations and exposure times during labor considered in this paper. Should it be found to decrease for lower concentrations, the oxytocin override exposure times corresponding to the indicated Pitocin concentrations during labor in Figure 4 would likewise decrease.

The factors discussed above in Subsections 8.1 to 8.12 may serve to reduce the time needed for given oxytocin override concentrations to cause desensitization of the OTRs in the fetal brain, as indicated in Figure 4, which may lead to offspring autism development.

## III. DISCUSSION OF EPIDEMIOLOGICAL STUDIES (SECTIONS 9 AND 10)

### 9. AN EXPLANATION FOR THE WEAK CORRELATION OBSERVED IN RECENT EPIDEMIOLOGICAL STUDIES

This section will attempt to clarify the finding of a weak correlation inferred for Pitocin use during labor and the subsequent development of autism in the offspring observed in recent epidemiological studies by Duke University in 2013 [1], Yale University in 2015 [2], and initially in the study by Harvard University in 2016 [3], mentioned in the Introduction. What is lacking in the attempts by these studies to interpret the findings of a weak correlation is the realization that there may be a desensitization, and hence; a down regulation threshold for the use of Pitocin during labor, below which no desensitization and subsequent down regulation will occur. This desensitization threshold is the product of the Pitocin concentration and the exposure time of OTRs in the fetal brain to the Pitocin equal to D = 1.8E6 (pg-min)/hr, which by the analysis in this paper can be related to parameters such as quantity of Pitocin used during labor, Pitocin infusion rate, maternal weight, length of time for labor and half-life for oxytocin removal in the maternal circulation that have been discussed in Sections 3, 4, 5, 6 and 7. Consequently, for Pitocin use below this threshold, there will be no OTR desensitization in the brain of the fetus, and hence; no correlation between Pitocin use during labor and the possible development of autism among the offspring. However, above this threshold, a correlation may exist.

From the analysis elsewhere in this paper and in a previous paper by this author [7], most labors will require much less Pitocin than the amount needed for an oxytocin override concentration to occur, or if it occurs, to reach the desensitization thresholds indicated in Figure 4. Hence, for most labors there will be no development of autism among the offspring that is attributable to OTR desensitization arising from the use of Pitocin, and subsequent down regulation. However, as indicated in Figure 4, for long labors with long Pitocin infusion times at high Pitocin infusion rates, a desensitization threshold may be reached and exceeded, resulting in the desensitization of OTR receptors in the brain of the fetus with subsequent down-regulation. It may be possible to establish correlations of long labors having long Pitocin infusion times with high Pitocin infusion levels with offspring autism development above this potential desensitization threshold.

The weak correlations indicated in the epidemiological studies may be explained in terms of Pitocin usage by combining no correlation for the majority of the cases of Pitocin use that are below a potential desensitization threshold with a much smaller number of cases having modest to strong correlations that are above this potential desensitization threshold. This combination may result in a weak correlation for the entire population.

A more exacting analysis may be done by examining the medical information concerning Pitocin usage, if it is available, that is associated with the cases examined in the epidemiological studies by Duke University, Yale University and Harvard University, where Pitocin use and offspring development of autism occur. These epidemiological studies did not examine the medical information associated with Pitocin use in the cases they examined to enable possible correlations of Pitocin use with offspring autism development. The possibility of such correlations indicates the necessity for these studies to be pursued with detailed epidemiological analysis.

Such examination of the medical information concerning Pitocin use associated with the epidemiological studies, if it is possible, may permit the determination of a possible correlation between Pitocin use and the subsequent offspring autism development. By examining the correlation, if it is found, it may be possible to identify an actual empirical desensitization threshold. Further analysis of the medical information on Pitocin use, if it is available, may permit the determination of risk factors for the development of autism correlated to quantity or units of Pitocin used, Pitocin infusion rate, length of time for labor and maternal weight, for cases above the empirical desensitization threshold that may be established. It may also be possible to establish a correlation between these factors and the severity of the autism in the offspring.

### 10. RE-ASSESSMENT OF HARVARD EPIDEMEOLOGICAL STUDY FINDINGS AND CONCLUSION

The approach indicated above should be seen in-light-of the findings of the recent epidemiological study by Harvard University of the occurrence of autism among the offspring of mothers who underwent labor induction with or without Pitocin in Sweden (A.S. Oberg, 2016) [3]. The initial data for the Harvard study suggested the possibility of a correlation between labor induction and the subsequent development of autism in offspring. However, the Harvard study used original statistical methods involving a family model to demonstrate that there was no correlation between the occurrence of autism during labor induction, and by inference, with the use of Pitocin, with a hazard ratio of 0.99 for a 95% confidence interval of (0.88-1.10) for their complete case category.

The analysis in this paper indicates that autism attributable to the use of Pitocin during labor would not occur during the time frames for most labor inductions, which averages 6 hours [5] and can take up to 12 hours or more for many women undergoing continuous Pitocin infusion [6]. Consequently; a correlation between offspring autism development and Pitocin usage would not exist. However, by the analysis in this paper, offspring autism development may be associated with high Pitocin infusion rates during long labor inductions with long Pitocin infusion times, especially; for smaller mothers with less efficient kidneys and livers for the removal of oxytocin. The number of cases would be relatively small and may not be reflected in the statistics the Harvard study presented. One factor influencing this is that the Harvard study considered labor induction as-a-whole, and not just continuous Pitocin infusion for labor induction. Harvard’s analysis could dilute the indication of the possible influence of continuous Pitocin infusion during labor induction on offspring autism development that may not be evident in the statistics they presented for their complete case category.

Taking these factors into account and using other data from the family model used in the Harvard study, namely; their sensitivity analysis for the category of first born of maternal cousins, which indicates a hazard ratio of 1.02 with a 95% confidence interval of (0.83-1.26), there may be an indication of a correlation between labor induction with Pitocin and offspring autism development. First births are generally longer [30], and this category in the Harvard findings, first born of maternal cousins, may more readily reflect the outcome of autism attributable to Pitocin usage indicated in Figure 4, where long labor inductions with long Pitocin infusion times using large quantities of Pitocin are a factor in OTR desensitization, which may lead to offspring autism development.

Consequently, there may be a higher percentage of cases of offspring autism development in the category of first births of maternal cousins which has a 1.02 hazard ratio, than in the complete case category which has a 0.99 hazard ratio. This may suggest a weak correlation between labor induction and offspring autism development. A stronger correlation may have been suggested in the Harvard study had continuous Pitocin infusion been considered separately for labor induction. Labor induction with Pitocin accounted for 63% of induced labors in the United States in 2016 [31], and the percentage of induced labors in Sweden in 2016, when the Harvard study was conducted in Sweden, is likely similar.

This hazard ratio of 1.02 may reflect the small incidence of labors with high levels of continuous Pitocin infusion for long labor inductions with long Pitocin infusion times necessary to achieve the oxytocin override levels indicated in Figure 4, that could result in OTR desensitization in the brain of the fetus and possible offspring autism development. Consequently, the conclusion of the Harvard study, that “the findings of this study provide no support for a causal association between induction of labor and offspring development of autism”, and the inference that follows, that continuous Pitocin infusion during labor induction has no causal association with offspring development of autism, may be in error.

## IV. RESEARCH CONSIDERATIONS (SECTION 11)

### 11. RESEARCH TO ADDRESS THE FINDINGS OF THIS PAPER

11.1 An analysis of the medical information for the cases of autism among offspring whose mothers received Pitocin during labor, if this medical information is available, may indicate a correlation between the occurrence of offspring autism development from mothers who had long labors with large quantities of Pitocin with high Pitocin infusion rates above an empirically determined desensitization threshold. The previous epidemiological studies by Duke University in North Carolina, Yale University in Denmark and initially by Harvard University in Sweden mentioned in the Introduction and in Sections 9 and 10, infer a weak correlation between Pitocin use during labor with the subsequent development of autism in some of the offspring. The acquisition of medical information, if possible, on the number of Units of Pitocin used, the rates of Pitocin infusion, the length of the labors and the weights of the mothers during labor, for the cases where the offspring developed autism may enable the determination of modest to strong correlations of Pitocin use with offspring autism development above an empirically determined desensitization threshold.

The information needed for this determination should be available in the medical records of the mothers in the cases in the epidemiological studies where Pitocin use and offspring autism development was found to occur. Obtaining this information for these cases, if it is possible, and performing the subsequent statistical analysis would shed light on the validity of the desensitization model presented in this paper, and even more so, may provide information for medical practitioners to evaluate the risk of autism in the offspring of the patients they treat with continuous Pitocin infusion during labor.

11.2 Such a correlation between the use of high levels of Pitocin use over long labors and offspring autism development, if it is found, would argue for a more circumspect use of Pitocin during labor, lowering the infusion rates where possible for a lower total use of Pitocin. Along these lines, it may be useful to revisit the application of pulsatile infusion of oxytocin for labor induction. Research into pulsatile infusion of oxytocin indicates that pulsatile infusion is more consistent with the natural pulsed secretion of oxytocin from the pituitary gland than continuous Pitocin infusion. This research has found that pulsatile infusion used 80% less oxytocin than continuous infusion [32,33], with the highest infusion rates of pulsatile infusion of oxytocin in one study 43% lower than those for continuous infusion of oxytocin [34]. Pulsatile infusion of oxytocin has been found to be as safe and effective as continuous oxytocin infusion for labor induction [32,33,34,35] and may mitigate the concerns raised in this paper regarding possible oxytocin override concentrations high enough to cause desensitization in the brain of the fetus.

Further research is needed to more fully evaluate the potential benefit of pulsatile infusion to determine if the pulses of oxytocin themselves may produce high oxytocin override concentrations for the duration of the pulse, and if so; what the effects may be on OTR desensitization in the brain of the fetus. Pulsatile infusion of Pitocin has been found to be less safe and less effective than continuous Pitocin infusion for labor augmentation [32,33]. Recent clinical trials for further study of pulsatile infusion of Pitocin have been proposed or undertaken [36,37]. If it is found to be suitable for general use in hospital settings, the application of pulsatile Pitocin infusion could be used as an alternative to, or in conjunction with continuous Pitocin infusion for labor induction. Its potential utility for the purposes of this paper may be in keeping Pitocin use below possible desensitization thresholds that may be found to exist by the analysis of the epidemiological studies called for in Subsection 11.1.

11.3 The extrapolation used in this paper for the presumed proportionality of the oxytocin override concentration C and OTR exposure time T involving the desensitization unit D [7], that was introduced in Section 5, arising from the constant value of the desensitization unit D = 1.8E6 (pg-min)/ml = CT, must be examined with further research into the desensitization of oxytocin receptors as a function of the indicated range of oxytocin concentrations and OTR exposure times in Figure 4. This research could be similar to the research in the study referenced in Section 5 on desensitization as a function of Pitocin concentration and Pitocin exposure time [16]. This research is necessary to validate more precisely the conclusions of this paper and achieve a more precise understanding of the role of high levels of Pitocin infusion during long labors with long Pitocin infusion times and the possible development of autism among the offspring. It should be noted that in a study of the effects of oxytocin on prairie voles, it was found that long-term low-dose administration of oxytocin may have caused possible desensitization with detrimental effects persisting long beyond treatments themselves [38]. This finding may argue for extending the desensitization threshold to lower levels of oxytocin concentrations than indicated in the literature [16] and may support the extrapolations indicated in this paper underlying the relationship between oxytocin override concentrations and desensitization thresholds indicated in Figure 4.

11.4 If a relationship can be found between the efficiencies of the kidneys and liver for removing oxytocin and the results of standard tests of kidney and liver function, it may be possible to determine the half-life of oxytocin in the maternal circulation during labor without the use of additional Pitocin, which could interfere with the labor itself. This would enable the identification of mothers potentially at risk for high concentrations of oxytocin arising from continuous Pitocin infusion during long labors as indicated in Figure 3, which could lead to OTR desensitization. Identifying these at-risk mothers may enable precautions to be taken that may circumvent the possibility of offspring autism development.

11.5 The mathematical model in this paper describing oxytocin override assumes an upper limit for placental oxytocinase degradation in the maternal circulation, which has not been established and which may change over the course of the labor itself and may differ among the placentas of different mothers producing the oxytocinase. Oxytocin override of placental oxytocinase degradation of oxytocin may be possible to study in animal models to evaluate the claims for OTR desensitization made in this study for high oxytocin infusion rates.

11.6 The production of increased concentrations of oxytocin in the cerebral spinal fluid of the fetus, that may result from the concentration of Pitocin entering the cerebral spinal fluid from the maternal circulation and binding with the OTRs in the magnocellular neurons, as discussed in Subsection 8.5, may be shown if the generation of oxytocin producing magnocellular neurons from adult stem cells from human beings can be done. This may permit in vitro research, showing how increasing the concentration of Pitocin in the fluid surrounding these magnocellular neurons may result in further increase in the production of oxytocin from these neurons by means of oxytocin’s function as a neurotransmitter and neuromodulator acting on them.

11.7 The data to establish the value of the desensitization unit D is taken from rat uterine explants, not human neurons. The generation of neurons from adult stem cells from human beings, if possible, may permit further studies of desensitization and down regulation of OTRs on neurons. It may also permit a more accurate determination of the value of the desensitization unit D for OTRs in the neurons of human beings and may permit a more accurate determination if D varies over a range of oxytocin concentrations and exposure times in human neurons. And, it may permit determination if OTR gene variation that is associated with autism is associated with greater vulnerability of these OTRs to desensitization.

## CONCLUSION

Although there are gaps in the scientific information needed to fully support the thesis presented in this paper, it is plausible that OTR desensitization in the brain of the fetus is possible during long labors with long Pitocin infusion times with high continuous Pitocin infusion rates, especially; for smaller mothers with less efficient kidneys and livers for removing oxytocin. In making this claim, this paper is more descriptive than prescriptive, and calls for consideration of the medical information, if available, in the epidemiological studies by Duke University in 2013, Yale University in 2015 and Harvard University in 2016, in which autism among offspring occurred along with the use of Pitocin by their mothers during labor.

The quantity of Pitocin used, the average infusion rate, the length of the labor and the weight of the mother at the time of birth, are factors that could affect a correlation between the use of Pitocin during labor and the occurrence of autism among the offspring. This paper asserts that this correlation, if found, may occur above an empirically determined desensitization threshold associated with the desensitization of the OTRs in the brain of the fetus.

The weak correlations observed in the epidemiological studies referenced in Section 10 may be explained as resulting from the combination of no correlation between the use of Pitocin during labor with autism for the majority of labors, which are below an empirically determined desensitization threshold that may be established for Pitocin use, and a strong correlation for the much smaller number of labors that are above this desensitization threshold.

The findings of the Harvard epidemiological study are re-interpreted, and the its conclusion is questioned to allow for the possibility of offspring autism development arising from the use of high infusion rates of Pitocin over long labor inductions with long Pitocin infusion times, especially for smaller mothers with less efficient kidneys and livers for the removal of oxytocin.

This paper establishes the relationship between continuous Pitocin infusion rate, half-life and the maternal weight with oxytocin concentration in the maternal circulation, indicating the significance of what may be the previously unknown influence of maternal oxytocin half-life on oxytocin concentration in the maternal circulation. The concepts introduced of oxytocin override and the limit of placental oxytocinase degradation of oxytocin may explain the possible buildup of oxytocin in the fetal circulation and the possible desensitization of OTRs in the brain of the fetus that may follow for long labors with long Pitocin infusion times with high Pitocin infusion rates.

The concern of this paper is to argue that a determination be made if there is a correlation between Pitocin use during long labors and offspring autism development from medical information, if it is available, that is associated with the epidemiological studies by Duke University, Yale University and Harvard University, while presenting a mathematical model that strongly alludes to the plausibility of such a determination. Limitations in the availability of scientific information that could support the mathematical model presented in this paper preclude a more accurate determination of the possible correlation between Pitocin use during labor and the development of autism among the offspring. However, the inferences drawn from the mathematical model presented in this paper are, that for long labors with long Pitocin infusion times at high Pitocin infusion rates for smaller mothers with higher than average half-lives for oxytocin removal from the maternal circulation, the offspring may be at greater risk for OTR desensitization and the possible development of autism. Higher desensitization levels may be shown to indicate a greater risk and severity of autism.

The possibility of the correlation between Pitocin use during labor and offspring development of autism discussed above, which may affect several hundred to one thousand or more offspring a year in the United States, should be motivation to warrant further investigation of the medical information, if it is available, that is associated with the epidemiological studies by Duke University, Yale University and Harvard University. This would allow for the possible development of risk factors for the potential onset of autism among the offspring that may call for a more circumspect use of Pitocin during long labors involving large quantities of Pitocin at high infusion rates. This may include the application of pulsatile oxytocin administration for labor induction, as well as consideration of alternative medical procedures during long labors in lieu of the use of large quantities of Pitocin at high infusion rates.

Declaration of Interest: None.

This research did not receive any specific grant from funding agencies in the public, commercial or not-for-profit sector.

